# The genomic basis of mood instability: identification of 46 loci in 363,705 UK Biobank participants, genetic correlation with psychiatric disorders, and association with gene expression and function

**DOI:** 10.1101/549931

**Authors:** Joey Ward, Elizabeth M. Tunbridge, Cynthia Sandor, Laura M. Lyall, Amy Ferguson, Rona J. Strawbridge, Donald M. Lyall, Breda Cullen, Nicholas Graham, Keira J.A. Johnston, Caleb Webber, Valentina Escott-Price, Michael O’Donovan, Jill P. Pell, Mark E.S. Bailey, Paul J. Harrison, Daniel J. Smith

**Affiliations:** Institute of Health and Wellbeing, University of Glasgow, Glasgow, UK; Department of Psychiatry, University of Oxford, UK; Oxford Health NHS Foundation Trust, Cardiff University, Cardiff, UK; UK Dementia Research Institute, Cardiff University, Cardiff, UK; Department of Medicine Solna, Karolinska Institute, Stockholm, Sweden; Department of Physiology, Anatomy and Genetics, Oxford, UK; University of Cardiff, Cardiff, UK; School of Life Sciences, College of Medical, Veterinary and Life Sciences, University of Glasgow, Glasgow, UK

## Abstract

Genome-wide association studies (GWAS) of psychiatric phenotypes have tended to focus on categorical diagnoses, but to understand the biology of mental illness it may be more useful to study traits which cut across traditional boundaries. Here we report the results of a GWAS of mood instability as a trait in a large population cohort (UK Biobank, n = 363,705). We also assess the clinical and biological relevance of the findings, including whether genetic associations show enrichment for nervous system pathways. Forty six unique loci associated with mood instability were identified with a SNP heritability estimate of 9%. Linkage Disequilibrium Score Regression (LDSR) analyses identified genetic correlations with Major Depressive Disorder (MDD), Bipolar Disorder (BD), Schizophrenia, anxiety and Post Traumatic Stress Disorder (PTSD). Gene-level and gene set analyses identified 244 significant genes and 6 enriched gene sets. Tissue expression analysis of the SNP-level data found enrichment in multiple brain regions, and eQTL analyses highlighted an inversion on chromosome 17 plus two brain-specific eQTLs. Additionally, we used a Phenotype Linkage Network (PLN) analysis and community analysis to assess for enrichment of nervous system gene sets using mouse orthologue databases. The PLN analysis found enrichment in nervous system PLNs for a community containing serotonin and melatonin receptors. In summary, this work has identified novel loci, tissues and gene sets contributing to mood instability. These findings may be relevant for the identification of novel trans-diagnostic drug targets and could help to inform future precision medicine innovations in mental health.

## Introduction

Mood instability is a subjective emotional state defined as rapid oscillations of intense affect, with difficulty regulating these oscillations and their behavioural consequences^1^. As a psychopathological phenotype, mood instability may be useful for psychiatric research within a Research Domain Classification (RDoC) framework^2^ because it is a symptom that occurs in several psychiatric disorders, particularly major depressive disorder (MDD) and bipolar disorder (BD). It is also present within general population samples, and is known to be associated with a range of adverse health outcomes^3^.

We recently identified four loci associated with mood instability within a subsample of the UK Biobank cohort (n = 113,968) and found genetic correlation with both MDD and schizophrenia^4^. Here, we report a significantly larger genome-wide association study (GWAS) of mood instability in the European ancestry subset of UK Biobank dataset (n = 363,705), using a BOLT-LMM approach to maximize statistical power. We also revisit the assessment of genetic correlations with psychiatric disorders, including the use of more recent GWAS outputs for MDD, schizophrenia and BD. Furthermore, we contextualize our findings in terms of affected tissues, eQTL analysis and Phenotype Linkage Network (PLN) analysis. PLN is a new methodology that harnesses the fact that variation in many complex traits results from perturbations of multiple molecular components within a smaller number of cellular pathways. These pathways can then be identified using gene network approaches.

## Methods

### UK Biobank sample

UK Biobank is a large cohort of over 500,000 United Kingdom residents, aged between 39 and 69 years^5^. UK Biobank was created to study the genetic, environmental and lifestyle factors that cause or prevent a range of morbidities in middle and older age. Baseline assessments occurred over a 4-year period, from 2006 to 2010, across 22 UK centres. These assessments covered a wide range of social, cognitive, lifestyle and physical health measures. Informed consent was obtained from all participants, and this study was conducted under generic approval from the NHS National Research Ethics Service (approval letter dated 13 May 2016, Ref 16/NW/0274) and under UK Biobank approvals for application #6553 ‘Genome-wide association studies of mental health’ (PI Daniel Smith).

### Genotyping, imputation and quality control

In March 2018 UK Biobank released genetic data for 487,409 individuals, genotyped using the Affymetrix UK BiLEVE Axiom or the Affymetrix UK Biobank Axiom arrays (Santa Clara, CA, USA) which contain over 95% common SNP content^6^. Pre-imputation quality control, imputation and post-imputation cleaning were conducted centrally by UK Biobank (described in the UK Biobank release documentation).

### Phenotyping

UK Biobank participants were asked as part of their baseline assessment: “Does your mood often go up and down?” Those who responded ‘yes’ to this question were defined as mood instability cases and those who responded ‘no’ were defined as controls. To minimise any impact of psychiatric disorders on observed genetic associations with mood instability, individuals reporting depression, bipolar disorder, schizophrenia, ‘nervous breakdown’, self-harm or suicide attempt (all from UK Biobank data field 20002), and those who reported taking psychotropic medications (data field 20003) were excluded from the analysis. Participants were also excluded if: their self-reported sex did not match their genetically determined sex; UK Biobank had determined them to have sex chromosome aneuploidy; they were considered by UK Biobank to be heterozygous outliers; they were missing over 10% of their genetic data; or they were not in the subset classified as British participants of European ancestry.

### Genetic association and heritability

Genetic association analysis was performed using BOLT-LMM^7, 8^. This approach makes use of a genetic relationship matrix (GRM) to control as robustly as possible for population structure without the need to adjust the model for principal components (PCs), while maximising power by avoiding the need to exclude related individuals. Additionally, BOLT-LMM builds an infinitesimal model including all directly genotyped SNPs simultaneously, thereby further increasing power compared to logistic regression approaches that test each SNP in turn. This ‘genotyped SNPs only’ model has the imputed SNPs tested against it allowing for the imputation score cut-off criterion to be substantially reduced and increases the number of SNPs available to test for association with the outcome. BOLT-LMM treats binary variables as a linear trait but is able to handle binary outcomes well when the sample size is large and when the number of cases and controls are evenly balanced, as is the case here.

Models were adjusted for age, sex and genotyping array. SNPs were filtered to remove those with MAF < 0.01, Hardy-Weinberg Equilibrium p < 1×10^-6^, or imputation quality score < 0.3. BOLT-LMM was also used to provide a heritability estimate and λ_GC_ estimate. A secondary analysis was also performed on a subsample of the cohort which excluded those used in the previous GWAS and anyone who was related to another participant.

Regional plots were made via LocusZoom v1.4^9^ as SNPs from the HRC reference panel were also imputed in the UK biobank genetic data release. We defined a locus as the region of containing a lead SNP and all other SNPs (r^2^ > 0.1) within a 5MB radius of the lead SNP. The LD was calculated using 10,000 unrelated Biobank participants who had passed the same genetic QC as those used for the GWAS.

The summary statistics were processed by FUMA^10^ (See URLs) for visualisation, MAGMA Gene Analysis, Gene-set Analysis and Tissue Expression Analysis^11^. The Gene-level Analysis operates by grouping p values for individual SNPs into a gene test statistic using the mean χ^2^ statistic for the gene whilst accounting for LD via the use of a European ancestry reference panel. The Gene-set Analysis groups genes according to MsigDB v6.1^12^, a collection of both curated gene sets and GO terms, and tests each set in turn against all the other sets. The Tissue Expression Analysis performs a one-sided test based on the correlation between tissue-specific gene expression profiles and trait-gene associations.

### Genetic correlations

Linkage Disequilibrium Score Regression (LDSR)^13^ was used to calculate genetic correlations with psychiatric disorders. The intercept was left unconstrained to allow for sample overlap. For the MDD^14^, BD^15^, schizophrenia^15^ and PTSD^16^ phenotypes, we used the most up-to-date GWAS summary statistics provided by the Psychiatric Genomics Consortium. Anxiety disorder summary statistics came from the Anxiety NeuroGenetics STudy (ANGST) Consortium ^17^.

### Tissue-specific expression and eQTL analysis

The lead SNP for each locus (unless otherwise noted) was assessed for *cis* effects on gene expression (eQTLs) in publicly available human dorsolateral prefrontal cortex RNASeq datasets using the Lieber Institute for Brain Development (LIBD) eQTL browser (See URLs). Each locus was initially examined in the LIIBD BrainSeq dataset (n = 738; See URLs); SNPs showing significant eQTLs were then assessed for replication in the Common Mind Consortium (CMC) dataset (n = 547; See URLs). Only eQTLs that reached a False Discovery Rate (FDR) corrected threshold of q ≤ 0.05 in both the LIBD and CMC datasets, and showed the same direction of effect in both, are reported. Tissue-specific expression patterns were assessed for implicated genes using the GTEx portal^18^. All q values quoted in the text are FDR corrected.

### Genetic principal component generation

Genetic principal components were created using plink 2^19^ using pca approx (with default settings) within the region between base positions 40,850,001 and 41,850,000 on chromosome 17 for the analysis of the inversion.

### Pathway analysis

PLN analysis builds on the fact that variation in complex traits results from perturbations of multiple molecular components within a smaller number of cellular pathways that can be identified using gene network approaches. No single dataset or data type can provide a complete picture of the functional association between genes but a recent method combines information from multiple data types by weighing functional similarities between genes according to their likelihood of influencing the same mammalian phenotype(s). This approach has a greater specificity and sensitivity than analyses using a single data type and other comparable integrative methods^20^. The PLN approach exploits phenotypic information from over seven thousand genes whose function has been experimentally perturbed in the mouse and evaluates the ability of different data types such as protein-protein interactions (PPI), co-expression (RNA or protein), and semantic similarity score based on literature or Gene Ontology (GO) annotations or pathway annotations (KEGG), to predict whether knockout of the orthologues of a given pair of human genes will yield similar phenotypes. By weighting those data types accordingly, they are integrated to generate a single combined measure of functional similarity between gene pairs. The resulting network of pairwise gene functional similarities is termed a phenotypic-linkage network (PLN)^20^.To increase the sensitivity and specificity to detect functional associations relevant for a specific disease/trait, it is possible to select only those mouse phenotypes that are relevant for a specific disorder in the data type weighting evaluation step^21^. Following this approach, we conducted a further analysis in which we re-weighted our generic PLN to be more sensitive to functional genomics data most informative to mood instability by considering only phenotypes within the over-arching mouse phenotype ontology (MPO) category *Nervous System* (MP:0003631). The PLN and nervous-PLN (NS-PLN) were built using the same 16 functional genomics datasets described by Honti et al^20^, with 64,640,972 and 49,656,123 weighted links respectively.

Following the approach described by Sandor et al., we identified ‘communities’ of densely interconnected groups of genes (including at least 20 genes) within each PLN and tested whether any communities were enriched in genes harboured by GWA/subGWA intervals. This test examines how many of these intervals harboured at least one gene belonging to a given Community as compared to randomly shifted intervals equal in gene number. This approach makes no prior hypothesis about the number or nature of genes within each GWA interval.

### Definition of GWA and subGWA intervals

The GWAS and subGWA intervals were defined by considering SNPs attaining an association p-value of 5×10^-8^ and 1×10^-6^ identified 6375 (GWA) and 9358 (sub-GWA) SNPs. We then identified the haplotypic block containing each of these of these SNPs using genotypes in the 1000 Genome Project and the python pipeline developed by Brent Pedersen (See URLs). We defined GWA/sub GWA intervals by identifying the most distant block on a chromosome within a region of 500Kb of the lead SNP. We then added an additional 300 kb on either side of the interval to include genes that may be regulated by regulatory variants with effects captured by the lead SNPs. For subGWA regions, we excluded those subGWA intervals harbouring genes present in GWA intervals.

## Results

### Demographics

In this GWAS sample of 363,705 individuals without a history of psychiatric disorder, 43.2% reported mood instability (n = 157,039) and the rest did not (n = 206,666). There was a higher proportion of females amongst the mood instability cases than in controls (55.4% versus 51.2% respectively), and the mean age of cases was lower than for controls (55.8 years versus 57.7 years).

### GWAS findings

We detected 46 loci across the genome with p < 5×10^-8^ (Figure 1 and Table S1) and an estimated SNP heritability (h^2^) of 0.09 (S.E. = 0.02). The heritability estimate has increased from the previous GWAS by approximately 2% (Previous h^2^ = 0.07, SE = 0.007). We attribute the increase in SE to the differing methodologies. The distribution of test statistics was consistent with a polygenic contribution to risk (λ_GC_ = 1.21; λ_1000_ = 1.001; LDSR intercept = 1.041; SE = 0.006). In addition, to help validate the four loci identified in the previous mood instability GWAS, we tested the top SNP from each locus in a sub-sample that excluded individuals in the previous smaller GWAS and those individuals related to another Biobank participant (n = 169,857). All four SNPs were associated with mood instability after Bonferroni correction (α < 0.0125, Table S2). We also note that the directions of effect were the same as for the previous GWAS findings.

**Figure 1.**
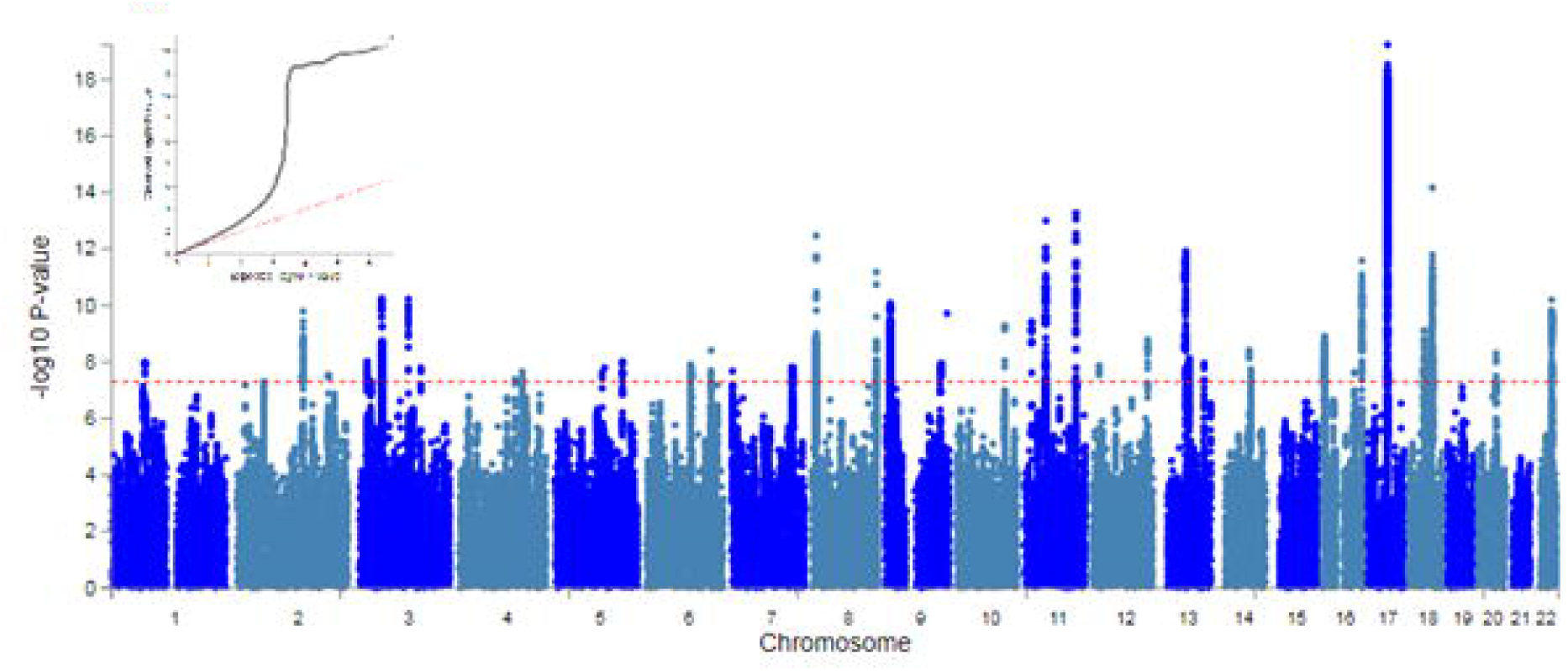
Manhattan and QQ plot of mood instability GWAS.

### Gene-level and Gene Set Analysis

244 significant genes were detected by MAGMA (Supplementary Table S3) and FUMA gene analysis. The Gene Set Analysis returned 6 enriched gene sets that met the threshold for significance after Bonferroni correction (Supplementary Table S4). Of these, 4 sets were related to brain development and differentiation of neurons, glial cells and astrocytes or neurogenesis. Other enriched sets included the Nikolsky breast cancer 16q24 amplicon genes and the prepulse inhibition gene sets.

### Tissue expression analysis

MAGMA tissue expression analysis identified 11 tissue categories, all of which were in the brain (Figure S1). Indeed, all sampled brain areas except substantia nigra showed enrichment (i.e., frontal and anterior cingulate cortex, basal ganglia, hippocampus, amygdala, hypothalamus and cerebellum). **Genetic correlations**

Genetic correlations were calculated between mood instability and five psychiatric phenotypes of interest (Table 1). All genetic correlations remained significant after FDR correction (q < 0.05). The largest correlations were with MDD (r_g_ = 0.74, q = 8.50*10^-157^) and anxiety (r_g_ = 0.64, q = 8.08*10^-6^). PTSD had a moderate correlation with mood instability (r_g_ = 0.32, q = 1.23*10^-2^) and both schizophrenia and bipolar disorder had weak but significant correlations (schizophrenia r_g_ = 0.14, q = 1.60*10^-5^, BD r_g_ = 0.09, q = 0.0012).

**Table 1.** Genetic correlations of mood instability with psychiatric phenotypes. r_g_ = genetic correlation with mood instability, S.E. = standard error of the genetic correlation, Z = the test statistic, p = the p value, q the False discovery rate corrected p value. MDD = major depressive disorder, PTSD = post-traumatic stress disorder.

### eQTL analysis

Nine of the GWAS loci showed significant eQTLs, many potentially with specific isoforms (summarised in Table S5 and presented in full in Supplementary Table S6). The strongest evidence of association with expression levels was for rs669915, an eQTL located within a region of strong linkage disequilibrium (LD) in chromosome 17q21 resulting from the existence of a 900kb inversion polymorphism that is common in European populations^22^. The extended region of LD across this portion of the chromosome makes it challenging to identify causal SNPs or the genes they regulate. The rs669915 eQTL was most strongly associated with expression of *LRRC37A4P* (LIBD dataset minimum q = 1.96 x 10^-9^; CMC dataset q = 3.99 x 10^-65^), an expressed pseudogene, but there are many alternative candidates for genes regulated by this SNP, including *MAPT* and *CRHR1*, for which it was also an eQTL. (Supplementary Table S5).

The chromosome 17q21 inversion polymorphism has itself been reported to affect the expression of genes in this region^23^. We therefore investigated whether rs669915 might ‘tag’ the expression effects mediated by the inversion polymorphism in our sample. Using the method of de Jong and colleagues, we constructed genetic principal components (GPCs) from SNPs within the region between base positions 40,850,001 and 41,850,000 on chromosome 17. A plot of the first two GPCs is shown in (Figure S2) and reveals three distinct clusters of individuals, each representing one of the three inversion polymorphism genotypes, H1/H1 (right-most cluster; n = 162,113), H1/H2 (middle cluster; n = 158,506) and H2/H2 (left-most cluster; n = 38,597). The H1 inversion allele had a population frequency of 0.32, far higher than the frequency reported by de Jong. In linear regression analyses, there was no association between mood instability phenotype and inversion genotype using a model of additive allelic effects (no. of H2 alleles) and adjusting for age, sex and genotyping array (p = 0.835).

### Nervous system PLN analyses (NS-PLN)

Amongst both GWA and subGWA gene sets, we found a disproportionate aggregation of genes within only one community, Community 26 within the NS-PLN (21 GWAS loci including at least one gene, q = 0.011; 25 “subGWAS “loci including at least one, q = 0.018) (**Fig 2A**). Examining the entire NS-PLN Community 26 gene, we found that it was significantly enriched in genes, whose unique 1:1 orthologues in the mouse when disrupted induce abnormities in synaptic transmission (Mouse Phenotype Ontology term MP:0003635; q = 2.77e^-118^, 75 genes expected vs 259 gene observed). However, we did not find evidence that the unique mouse orthologues of mood instability GWA and subGWA genes that belonged to Community 26 were enriched for any particular mouse phenotype. Nonetheless, we found that the 37 and 35 GWA and subGWA genes present in the Community 26 were highly functionally connected with other Community 26 genes annotated with abnormal synaptic transmission phenotype term (Figure 2B).

**Figure 2.**
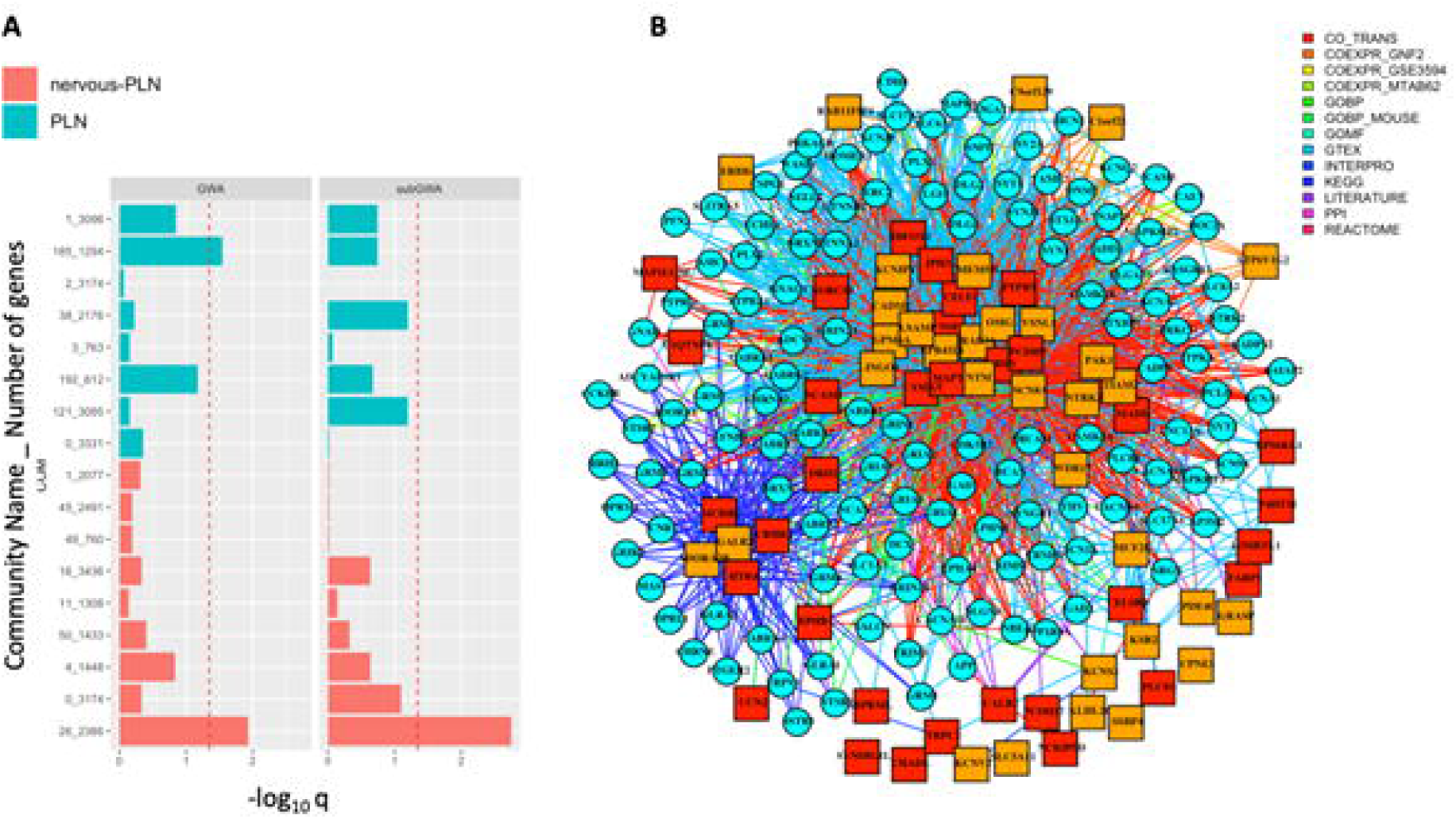
Different Mood associated genetic risk variants converge in a nervous specific gene network. (A) Enrichments of gene functional communities from a generic PLN and from a Nervous-System (NS) PLN within Mood-GWA and subGWA loci (see Methods). The Community ID is given first in the descriptor followed by the number of genes within that community. Only communities formed from over 20 genes are shown. (B) Gene subnetwork of Community 26 from NS-PLN showing functional associations between genes residing in Mood-associated GWA (red squares) and subGWA (orange squares) intervals and genes whose unique mouse orthologues are annotated with *abnormal synaptic transmission phenotype* (cyan squares). To increase clarity, only genes with *abnormal synaptic transmission phenotype* annotation with at least three functional links to genes residing in GWA and subGWA regions are shown. The colour of the link connecting two genes indicates the strongest information source supporting the functional association.

## Discussion

### Main findings

These analyses represent the largest genetic study of mood instability to date. Forty six unique loci associated with mood instability were identified, with a heritability estimate of 9%. Our findings confirm the four loci identified in our initial GWAS on the UK Biobank interim data release^4^ and are further validated by tissue expression analyses (enrichment for 11 brain regions) and pathway analyses (6 enrichment pathways, 4 of which relate to the development and differentiation of neurons). The large number of individuals in this study also provided substantial power to detect genetic correlations with psychiatric traits via LDSR. All five psychiatric traits assessed had a significant genetic correlation with mood instability. Some of these correlations were strong (particularly for MDD and anxiety) but others were weaker than expected. For example, the genetic correlation between mood instability and BD was only 9%, perhaps suggesting that the mood instability phenotype in this study differs from the affective instability that is a core feature of BD.

### Biology of mood instability

Loci associated with mood instability included genes that are involved across a variety of biochemical pathways, as well as brain development and function. For example, several gene products localised to the synapse. *PLCL1* and *PLCL2* are involved in GABA signalling^24^ and melatonin signalling respectively, and *RAPSN* assists in anchoring nicotinic acetylcholine receptors at synaptic sites^25^. *PLCL1* has already been identified in a GWAS of schizophrenia^26^ and *PLCL2* has been shown to be upregulated in bipolar disorder^27^. Additionally, we identified *CALB2* which has many biological functions, including a role in modulating neuronal excitability^28^. Both *DCC* (identified in the previous mood instability GWAS) and *BSN* facilitate the release of neurotransmitters within the active zone of some axons^29^. *BSN* has also been shown to be associated with schizoaffective disorder via GABA signalling^30^. *FARP1* promotes dendritic growth^31^ and, although it has so far not been directly linked to psychiatric disorders, it has been shown to regulate dendritic complexity^32^ (reduced dendritic complexity is a recognised feature of schizophrenia^33^).

We identified several developmental genes, including *NEGR1*^34^, *RARB*^35^ and *EPHB1*^36^, and transcription factors such as *HIVEP2* (loss of function of which causes intellectual disability^37^) and *TCF4* (previously associated with schizophrenia^38^). *NEGR1* was identified by 23andMe within their GWAS of MDD^39^ and increased levels of NEGR1 protein in spinal fluid have been identified in both MDD and BD^40^. *RARB* is involved in retinoic acid synthesis pathways that have been associated depressive symptoms in mice^41^ and has also been found to have increased expression in patients with schizophrenia^42^. The methylation state of the *EPHB1* gene has been linked to MDD^43^ and SNP-based analyses have identified association between *EPHB1*’s and symptoms of schizophrenia^44^.

We also found association with several genes involved in mitochondrial energy production, such as *NDUFAF3, NDUFS3, PTPMT1, KBTBD4* and *MTCH2*, suggesting that part of the physiology of mood instability may relate to energy dysregulation.

In addition to protein coding genes, several loci were identified in regions containing non-coding protein sequences such as *AC019330.1, AC133680.1, RP11-6N13.1* and *RP11-436d23.1*. Additionally, eQTL analyses identified three more possible non-coding genes (*RP11-481A20.10, RP11-481A20.11* and *FAM85B*) suggesting a possible RNA interference or post-transcriptional regulation basis to mood instability.

Furthermore, the eQTL analyses highlighted the 17q21 inversion. Our principal component analysis of this region did not detect a significant association leading us to conclude that it is the SNPs in the region (not the inversion itself) driving the association. It is possible that lead SNPs may tag, enhancer RNA or eRNA which we were unable to detect here (LIBD data was generated using poly-A RNA and so targets messenger RNA). However, our findings are consistent with a recent report implicating dopamine neuron-enriched enhancer activity in this region in several dopamine-related psychiatric and neurological conditions^45^.

Genes within regions associated with mood instability were functionally associated with synaptic transmission, a key pathway for psychiatric disorders, albeit this functional association was only detectable after focussing our gene network towards data types most informative for mammalian nervous system phenotypes. Among the genes lying within associated loci that contribute to this functional association are several interesting candidate genes. *HTR4* is a member of the family of serotonin receptors and implicated in depression and its treatment^46, 47^. *MCHR1*, melanin concentrating hormone receptor 1, is a G protein-coupled which binds melanin-concentrating hormone. *MCHR1* can inhibit cAMP accumulation and stimulate intracellular calcium flux, and may be involved in the neuronal regulation of food consumption^46^ and the locus showed association with schizophrenia in a Danish sample^49^.

### Strengths and Limitations

As noted above, this is the largest GWAS of a mood instability phenotype to date and has successfully identified new loci, eQTLs, tissues, genetic correlations and gene network enrichments. However, there are several limitations, most notably the use of a single question to define mood instability, and the lack of objective verification of this phenotype. There are more detailed suggested measurement scales for mood instability, such as that developed by Chaturvedi and colleagues^50^. In the future, the use of these more comprehensive assessments in large samples may provide some clarification of our findings, for example the lack of strong genetic correlation between mood instability and BD. Nevertheless, the single question approach to mood instability has been widely used, and shown to identify robust associations with a range of health outcomes and disorders^1, 3^. Similarly, exclusions for psychiatric disorder were based on self-report.

We validated the top hits of the previous GWAS however the cohort used was not truly independent (it was also part of the UK Biobank cohort). It would be of interest to replicate the 46 loci identified here should sufficiently large independent replication cohorts become available in the future. As well as replication of the loci, further analysis of sex specific differences would be of interest because mood instability was more common in females than males. Although this difference was relatively small, our reported analyses were adjusted for sex and these differences are similar to those reported elsewhere^51^.

It is also important to note that direct links between genetic risk loci and network constituents in the PLN analysis will have to await the release of more completely annotated gene databases. The incompleteness of phenotypic annotations is likely to explain why the genes identified in the PLN analysis don’t have corresponding organismal or physiological phenotypes, but the fact that there were strong functional associations between the genes in the network we detected and mouse orthologues that have the synaptic transmission phenotype annotation suggests that the mood instability genes will also reveal this phenotype when more completely annotated databases become available.

Finally, we note the large difference in frequencies of the inversion polymorphism on chromosome 17 from that reported by De Jong^23^. This difference could be due to the populations sampled to estimate the frequency or over representation in those who joined UK Biobank. It is however important to note that the inversion itself would be likely to contribute only a small proportion of the mood instability phenotype, such that even larger sample sizes than were used here would be needed to detect a correlation where one exists..

## Conclusion

In summary, with a tripling in sample size from the previous GWAS, we identified substantially more associations with mood instability in the UK Biobank cohort^4^. Future analyses of the precise roles that these associations play in the clinical expression of mood instability will be relevant for the wide range of psychiatric phenotypes in which mood instability occurs. We have also been able to more confidently place these GWAS findings within relevant biological contexts and some of the loci and pathways identified may represent candidates for future novel drug development.

Our findings are also of interest in the context of precision medicine innovations for mental health. It is possible that polygenic risk scores derived from this work could be applied to clinical populations to conduct pharmacogenomics studies and to inform patient stratification approaches. Overall, we hope that our findings will stimulate further research on the biology and treatment of mood instability across a range of mood and psychotic disorders.

## URLs

FUMA - http://fuma.ctglab.nl/

Python pipeline developed by Brent Pedersen - https://gist.github.com/brentp/5050522

LIBD website - http://eqtl.brainseq.org/

LIBD eQTL Browser phase 1 - http://eqtl.brainseq.org/phase1/eqtl/

CommonMind Consortium public–private partnership http://commonmind.org/WP

## Acknowledgements

JW is supported by the JMAS Sim Fellowship for depression research from the Royal College of Physicians of Edinburgh (173558). AF is supported by an MRC Doctoral Training Programme Studentship at the University of Glasgow (MR/K501335/1). RJS is supported by a UKRI Innovation-HDR-UK Fellowship (MR/S003061/1). KJAJ is supported by an MRC Doctoral Training Programme Studentship at the Universities of Glasgow and Edinburgh. DJS acknowledges the support of a Lister Prize Fellowship (173096) and the MRC Mental Health Data Pathfinder Award (MC_PC_17217). EMT and PJH are supported by the Oxford Health NIHR Biomedical Research Centre. The views expressed are those of the authors and not necessarily those of the National Health Service, NIHR or the Department of Health. Data were generated as part of the CommonMind Consortium supported by funding from Takeda Pharmaceuticals Company Limited, F. Hoffman-La Roche Ltd and NIH grants R01MH085542, R01MH093725, P50MH066392, P50MH080405, R01MH097276, RO1-MH-075916, P50M096891, P50MH084053S1, R37MH057881 and R37MH057881S1, HHSN271201300031C, AG02219, AG05138 and MH06692. Brain tissue for the study was obtained from the following brain bank collections: the Mount Sinai NIH Brain and Tissue Repository, the University of Pennsylvania Alzheimer’s Disease Core Center, the University of Pittsburgh NeuroBioBank and Brain and Tissue Repositories and the NIMH Human Brain Collection Core. CMC Leadership: Pamela Sklar, Joseph Buxbaum (Icahn School of Medicine at Mount Sinai), Bernie Devlin, David Lewis (University of Pittsburgh), Raquel Gur, Chang-Gyu Hahn (University of Pennsylvania), Keisuke Hirai, Hiroyoshi Toyoshiba (Takeda Pharmaceuticals Company Limited), Enrico Domenici, Laurent Essioux (F. Hoffman-La Roche Ltd), Lara Mangravite, Mette Peters (Sage Bionetworks), Thomas Lehner, Barbara Lipska (NIMH).

We thank all participants in the UK Biobank study. UK Biobank was established by the Wellcome Trust, Medical Research Council, Department of Health, Scottish Government and Northwest Regional Development Agency. UK Biobank has also had funding from the Welsh Assembly Government and the British Heart Foundation. Data collection was funded by UK Biobank.

The summary statistics of the GWAS are available upon request by contacting the corresponding author.

---

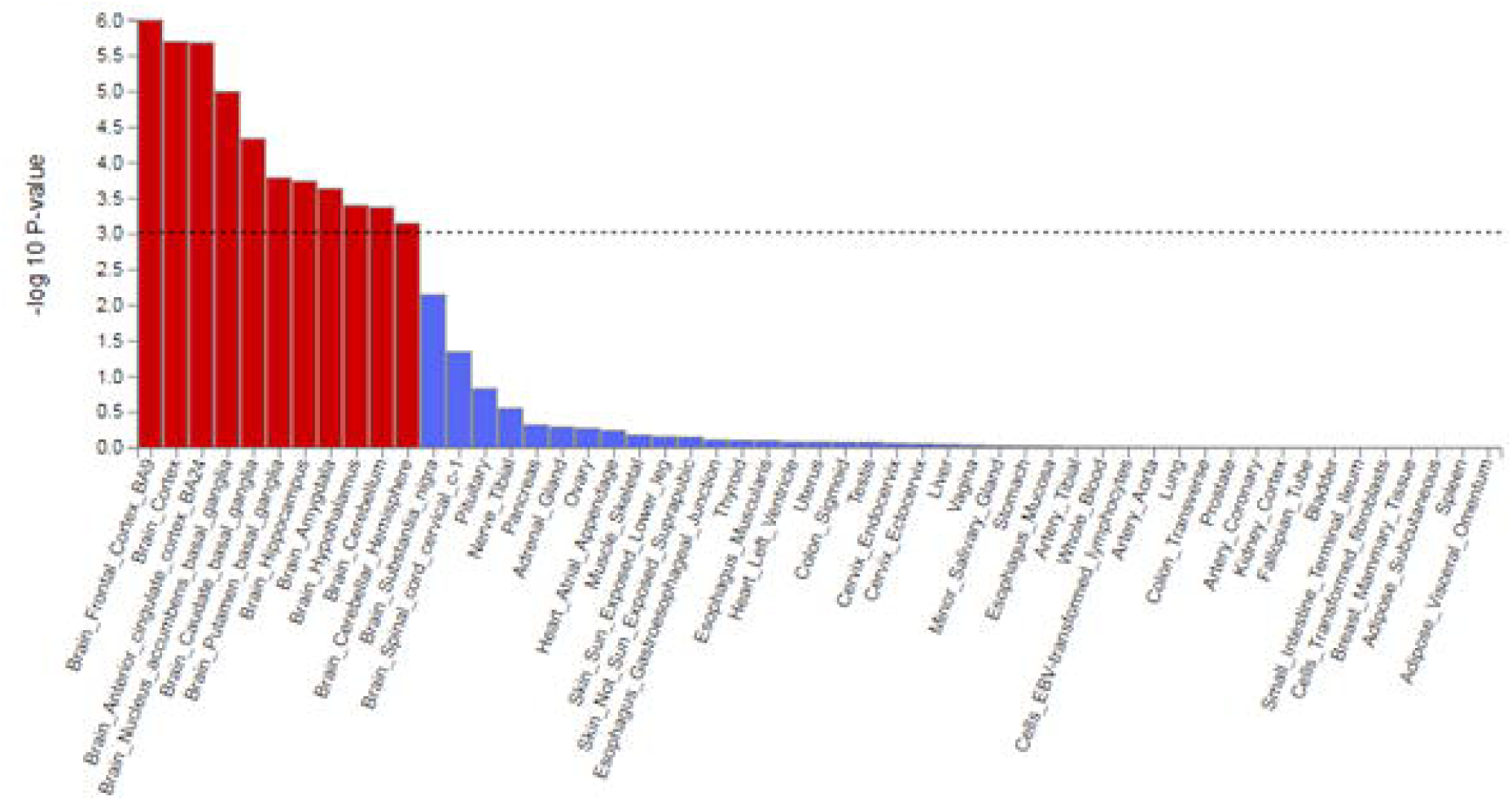

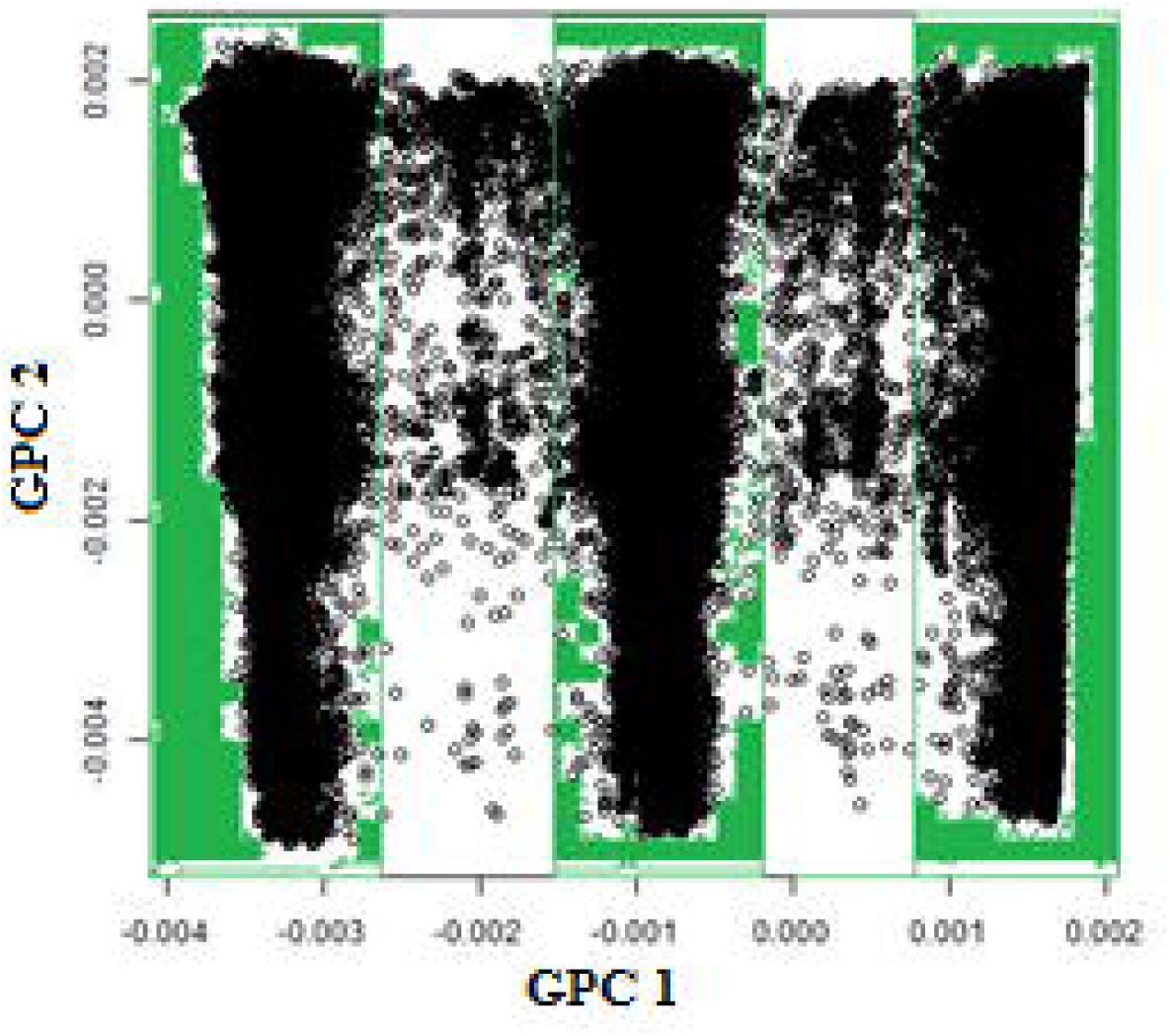

## References

1. Marwaha S, He Z, Broome M, Singh SP, Scott J, Eyden J et al. How is affective instability defined and measured? A systematic review. Psychol Med 2014; 44(9): 1793–1808.

2. Cuthbert BN, Insel TR. Toward the future of psychiatric diagnosis: the seven pillars of RDoC. BMC Med 2013; 11: 126.

3. Broome MR, Saunders KEA, Harrison PJ, Marwaha S. Mood instability: significance, definition and measurement. The British journal of psychiatry: the journal of mental science 2015; 207(4): 283–285.

4. Ward J, Strawbridge RJ, Bailey MES, Graham N, Ferguson A, Lyall DM et al. Genome-wide analysis in UK Biobank identifies four loci associated with mood instability and genetic correlation with major depressive disorder, anxiety disorder and schizophrenia. Translational Psychiatry 2017; 7(11): 1264.

5. Sudlow C, Gallacher J, Allen N, Beral V, Burton P, Danesh J et al. UK biobank: an open access resource for identifying the causes of a wide range of complex diseases of middle and old age. PLoS Med 2015; 12(3): e1001779.

6. Biobank U. Genotyping of 500,000 UK Biobank participants. Description of sample processing workflow and preparation of DNA for genotyping. 2015;11 September 2015.

7. Loh PR, Tucker G, Bulik-Sullivan BK, Vilhjalmsson BJ, Finucane HK, Salem RM et al. Efficient Bayesian mixed-model analysis increases association power in large cohorts. Nat Genet 2015; 47(3): 284–290.

8. Loh P-R, Kichaev G, Gazal S, Schoech AP, Price AL. Mixed-model association for biobank-scale datasets. Nature Genetics 2018.

9. Watanabe K, Taskesen E, van Bochoven A, Posthuma D. Functional mapping and annotation of genetic associations with FUMA. Nature Communications 2017; 8(1): 1826.

10. de Leeuw CA, Mooij JM, Heskes T, Posthuma D. MAGMA: Generalized Gene-Set Analysis of GWAS Data. PLOS Computational Biology 2015; 11(4): e1004219.

11. Subramanian A, Tamayo P, Mootha VK, Mukherjee S, Ebert BL, Gillette MA et al. Gene set enrichment analysis: A knowledge-based approach for interpreting genome-wide expression profiles. Proceedings of the National Academy of Sciences 2005; 102(43): 15545.

12. Pruim RJ, Welch RP, Sanna S, Teslovich TM, Chines PS, Gliedt TP et al. LocusZoom: regional visualization of genome-wide association scan results. Bioinformatics (Oxford, England) 2010; 26(18): 2336–2337.

13. Bulik-Sullivan BK, Loh P-R, Finucane HK, Ripke S, Yang J, Schizophrenia Working Group of the Psychiatric Genomics C et al. LD Score regression distinguishes confounding from polygenicity in genome-wide association studies. Nat Genet 2015; 47(3): 291–295.

14. Wray NR, Ripke S, Mattheisen M, Trzaskowski M, Byrne EM, Abdellaoui A et al. Genome-wide association analyses identify 44 risk variants and refine the genetic architecture of major depression. Nat Genet 2018; 50(5): 668–681.

15. Genomic Dissection of Bipolar Disorder and Schizophrenia, Including 28 Subphenotypes. Cell 2018; 173(7): 1705-1715.e1716.

16. Duncan LE, Ratanatharathorn A, Aiello AE, Almli LM, Amstadter AB, Ashley-Koch AE et al. Largest GWAS of PTSD (N=20 070) yields genetic overlap with schizophrenia and sex differences in heritability. Mol Psychiatry 2018; 23(3): 666–673.

17. Otowa T, Hek K, Lee M, Byrne EM, Mirza SS, Nivard MG et al. Meta-analysis of genome-wide association studies of anxiety disorders. Molecular psychiatry 2016; 21(10): 1391–1399.

18. Consortium GT. The Genotype-Tissue Expression (GTEx) project. Nature genetics 2013; 45(6): 580–585.

19. Purcell S, Neale B, Todd-Brown K, Thomas L, Ferreira Manuel A R, Bender D et al. PLINK: A Tool Set for Whole-Genome Association and Population-Based Linkage Analyses. American Journal of Human Genetics 2007; 81(3): 559–575.

20. Honti F, Meader S, Webber C. Unbiased functional clustering of gene variants with a phenotypic-linkage network. PLoS Comput Biol 2014; 10(8): e1003815.

21. Sandor C, Beer NL, Webber C. Diverse type 2 diabetes genetic risk factors functionally converge in a phenotype-focused gene network. PLoS Comput Biol 2017; 13(10): e1005816.

22. Stefansson H, Helgason A, Thorleifsson G, Steinthorsdottir V, Masson G, Barnard J et al. A common inversion under selection in Europeans. Nature Genetics 2005; 37: 129.

23. de Jong S, Chepelev I, Janson E, Strengman E, van den Berg LH, Veldink JH et al. Common inversion polymorphism at 17q21.31 affects expression of multiple genes in tissue-specific manner. BMC Genomics 2012; 13: 458.

24. Kanematsu T, Jang I-S, Yamaguchi T, Nagahama H, Yoshimura K, Hidaka K et al. Role of the PLC-related, catalytically inactive protein p130 in GABA(A) receptor function. The EMBO journal 2002; 21(5): 1004–1011.

25. Muller JS, Baumeister SK, Rasic VM, Krause S, Todorovic S, Kugler K et al. Impaired receptor clustering in congenital myasthenic syndrome with novel RAPSN mutations. Neurology 2006; 67(7): 1159–1164.

26. Schizophrenia Working Group of the Psychiatric Genomics C. Biological insights from 108 schizophrenia-associated genetic loci. Nature 2014; 511(7510): 421–427.

27. Nakatani N, Hattori E, Ohnishi T, Dean B, Iwayama Y, Matsumoto I et al. Genome-wide expression analysis detects eight genes with robust alterations specific to bipolar I disorder: relevance to neuronal network perturbation. Human Molecular Genetics 2006; 15(12): 1949–1962.

28. Camp AJ, Wijesinghe R. Calretinin: modulator of neuronal excitability. Int J Biochem Cell Biol 2009; 41(11): 2118–2121.

29. Winter C, tom Dieck S, Boeckers TM, Bockmann J, Kampf U, Sanmarti-Vila L et al. The presynaptic cytomatrix protein Bassoon: sequence and chromosomal localization of the human BSN gene. Genomics 1999; 57(3): 389–397.

30. Hamshere ML, Green EK, Jones IR, Jones L, Moskvina V, Kirov G et al. Genetic utility of broadly defined bipolar schizoaffective disorder as a diagnostic concept. Br J Psychiatry 2009; 195(1): 23–29.

31. Zhuang B, Su YS, Sockanathan S. FARP1 promotes the dendritic growth of spinal motor neuron subtypes through transmembrane Semaphorin6A and PlexinA4 signaling. Neuron 2009; 61(3): 359–372.

32. Cheadle L, Biederer T. Activity-Dependent Regulation of Dendritic Complexity by Semaphorin 3A through Farp1. The Journal of Neuroscience 2014; 34(23): 7999.

33. Broadbelt K, Byne W, Jones LB. Evidence for a decrease in basilar dendrites of pyramidal cells in schizophrenic medial prefrontal cortex. Schizophrenia Research 2002; 58(1): 75–81.

34. Takita J, Chen Y, Okubo J, Sanada M, Adachi M, Ohki K et al. Aberrations of NEGR1 on 1p31 and MYEOV on 11q13 in neuroblastoma. Cancer Sci 2011; 102(9): 1645–1650.

35. van der Wees J, Schilthuis JG, Koster CH, Diesveld-Schipper H, Folkers GE, van der Saag PT et al. Inhibition of retinoic acid receptor-mediated signalling alters positional identity in the developing hindbrain. Development 1998; 125(3): 545–556.

36. Wilkinson DG. Multiple roles of EPH receptors and ephrins in neural development. Nat Rev Neurosci 2001; 2(3): 155–164.

37. Srivastava S, Engels H, Schanze I, Cremer K, Wieland T, Menzel M et al. Loss-of-function variants in HIVEP2 are a cause of intellectual disability. European Journal of Human Genetics 2016; 24(4): 556–561.

38. Williams HJ, Moskvina V, Smith RL, Dwyer S, Russo G, Owen MJ et al. Association between TCF4 and schizophrenia does not exert its effect by common nonsynonymous variation or by influencing cis-acting regulation of mRNA expression in adult human brain. Am J Med Genet B Neuropsychiatr Genet 2011; 156b(7): 781–784.

39. Hyde CL, Nagle MW, Tian C, Chen X, Paciga SA, Wendland JR et al. Identification of 15 genetic loci associated with risk of major depression in individuals of European descent. Nature genetics 2016; 48(9): 1031–1036.

40. Maccarrone G, Ditzen C, Yassouridis A, Rewerts C, Uhr M, Uhlen M et al. Psychiatric patient stratification using biosignatures based on cerebrospinal fluid protein expression clusters. Journal of Psychiatric Research 2013; 47(11): 1572–1580.

41. O’Reilly KC, Shumake J, Gonzalez-Lima F, Lane MA, Bailey SJ. Chronic Administration of 13-Cis-Retinoic Acid Increases Depression-Related Behavior in Mice. Neuropsychopharmacology 2006; 31: 1919.

42. Tsai SY, Catts VS, Fullerton JM, Corley SM, Fillman SG, Weickert CS. Nuclear Receptors and Neuroinflammation in Schizophrenia. Molecular Neuropsychiatry 2017; 3(4): 181–191.

43. Davies MN, Krause L, Bell JT, Gao F, Ward KJ, Wu H et al. Hypermethylation in the ZBTB20 gene is associated with major depressive disorder. Genome Biology 2014; 15(4): R56–R56.

44. Su L, Ling W, Jiang J, Hu J, Fan J, Guo X et al. Association of EPHB1 rs11918092 and EFNB2 rs9520087 with psychopathological symptoms of schizophrenia in Chinese Zhuang and Han populations. Asia Pac Psychiatry 2016; 8(4): 306–308.

45. Dong X, Liao Z, Gritsch D, Hadzhiev Y, Bai Y, Locascio JJ et al. Enhancers active in dopamine neurons are a primary link between genetic variation and neuropsychiatric disease. Nature Neuroscience 2018; 21(10): 1482–1492.

46. Fontaine-Bisson B, Thorburn J, Gregory A, Zhang H, Sun G. Melanin-concentrating hormone receptor 1 polymorphisms are associated with components of energy balance in the Complex Diseases in the Newfoundland Population: Environment and Genetics (CODING) study. Am J Clin Nutr 2014; 99(2): 384–391.

47. Demontis D, Nyegaard M, Christensen JH, Severinsen J, Hedemand A, Hansen T et al. The gene encoding the melanin-concentrating hormone receptor 1 is associated with schizophrenia in a Danish case-control sample. Psychiatr Genet 2012; 22(2): 62–69.

